# Chronic delivery of buprenorphine during abstinence decreases incubation of heroin seeking and neuronal activation in medial prefrontal cortex and striatum in male and female rats

**DOI:** 10.1101/2025.09.20.677438

**Authors:** Jennifer M. Bossert, Kiera E. Caldwell, Hannah Bonbrest, Rohan Patil, Mona S. Pishgar, Rajtarun Madangopal, Huiling Wang, Yavin Shaham

**Affiliations:** Behavioral Neuroscience Branch, IRP/NIDA/NIH, Baltimore, MD, U.S.A.

## Abstract

**Rationale and Objective:** Buprenorphine is an FDA-approved medication for opioid addiction, but the brain regions underlying its therapeutic effects remain unknown. We previously found that chronic buprenorphine treatment decreases several relapse-related behaviors in rats. Here, we tested whether chronic buprenorphine decreases the time-dependent increase (incubation) in heroin seeking during abstinence. We also used the activity marker Fos to test whether buprenorphine’s inhibition of incubation of heroin seeking is associated with decreased activation of cortical and striatal regions.

**Methods:** We trained *Oprm1-Cre* rats and their wild-type littermates to self-administer heroin for 12 days (6 h/day). On abstinence Day 1, we tested for heroin seeking under extinction conditions. On Day 14, we implanted osmotic minipumps containing saline or buprenorphine (6 mg/kg/day). On abstinence Days 21-22, we tested the rats (or not tested, baseline condition) for incubation of heroin seeking, after which brains were collected for Fos immunohistochemistry.

**Results:** *Oprm1-Cre* rats did not differ from wild-type littermates in heroin self-administration or ‘non-incubated’ heroin seeking on abstinence Day 1. In both genotypes, chronic buprenorphine decreased incubated heroin seeking on Days 21-22. Buprenorphine also decreased incubation-related neuronal activation in several cortical areas (anterior cingulate and dorsal peduncular, but not prelimbic, infralimbic, or orbitofrontal cortex) and striatal areas (dorsolateral and dorsomedial striosomes and nucleus accumbens core, but not shell).

**Conclusions:** Chronic buprenorphine decreased incubation of heroin seeking, supporting the predictive validity of the rat model. This effect was associated with decreased neuronal activity in specific subregions of the medial prefrontal cortex and striatum.

## Introduction

Relapse after prolonged abstinence is a defining feature of opioid addiction (Hunt et al. 1971; Sinha 2011) and is often triggered by exposure to drug-associated cues and contexts that provoke drug craving (O’Brien et al. 1992). In laboratory rats, “incubation of drug craving” refers to the time-dependent increase in drug seeking during abstinence (Chow et al. 2025; Grimm et al. 2001). We and others have shown that opioid seeking progressively increases, or incubates, during abstinence from heroin (Shalev et al. 2001), fentanyl (Gyawali et al. 2020), and oxycodone (Blackwood et al. 2019; Fredriksson et al. 2020) self-administration.

Buprenorphine, a partial μ-opioid receptor (MOR) agonist and κ-opioid receptor (KOR) antagonist, is an effective FDA-approved treatment for opioid addiction (Jasinski et al. 1978; Kakko et al. 2003). In a previous study using a rat model of opioid maintenance therapy (Leri et al. 2004; Shaham et al. 1996; Sorge et al. 2005), we found that buprenorphine decreased several relapse-related behaviors in outbred Sprague–Dawley rats of both sexes (Bossert et al. 2020). These behaviors included extinction responding, context-induced reinstatement, and reacquisition of oxycodone self-administration. Here, we examined whether chronic buprenorphine also decreases the incubation of heroin seeking in male and female *Oprm1-Cre* rats (Bossert et al. 2023) and their wild-type littermates.

The brain regions underlying the therapeutic effects of buprenorphine in humans and in the rat opioid maintenance model remain unknown. To begin addressing this question, we tested whether buprenorphine-induced inhibition of incubation of heroin seeking is associated with decreased activation (measured using the activity marker Fos (Cruz et al. 2013)) in cortical and striatal regions that express MOR (Mansour et al. 1988). These include medial prefrontal cortex (mPFC: anterior cingulate, prelimbic, infralimbic, and dorsal peduncular), orbitofrontal cortex (OFC: lateral and ventral), and striatum (dorsolateral, dorsomedial, nucleus accumbens [NAc] core, and NAc shell).

Regarding the mPFC, while a causal role in the incubation of opioid seeking has not been established (Chow et al. 2025), extensive evidence implicates different mPFC subregions in cue- and context-induced reinstatement of heroin seeking (Bossert et al. 2011; Reiner et al. 2019; Rogers et al. 2008) and in the incubation of cocaine seeking (Chiu et al. 2021; Shin et al. 2018; Szumlinski et al. 2019). For the OFC, several studies using causal manipulations (gene knockdown, Daun02 inactivation, chemogenetics) have demonstrated its role in the incubation of opioid seeking (heroin and oxycodone) (Altshuler et al. 2021; Fanous et al. 2012; Lin et al. 2024; Olaniran et al. 2023; Zanda et al. 2023). For the striatum, causal evidence supports roles for both dorsal and ventral subregions in incubation of opioid seeking (Lin et al. 2024; Sun et al. 2015; Wong et al. 2022; Xu et al. 2021).

Within the dorsal striatum, we also examined Fos expression in striosomes (patches) and the surrounding matrix, given their distinct roles in motivated behavior (Friedman et al. 2017; Friedman et al. 2015). Striosomes were identified with an anti-MOR antibody, as this compartment exhibits high MOR expression (Graybiel 1990; Graybiel and Ragsdale 1978; Herkenham and Pert 1981; Pert et al. 1976).

Finally, we used newly developed *Oprm1-Cre* rats (Bossert et al. 2023) and their wild-type littermates. Our initial goal was to co-localize Fos and nuclear Cre expression in *Oprm1-Cre* rats to determine whether buprenorphine selectively decreases activity in *Oprm1*-expressing cells. However, despite testing three anti-Cre antibodies raised in rabbit, mouse, and guinea pig, we were unable to obtain specific labeling.

## Methods

### Subjects

We used 56 rats (29 *Oprm1*-Cre heterozygotes [9 males and 20 females] and 27 wildtype littermates [12 males and 13 females]) from the NIDA IRP Transgenic Breeding Core. The rats weighed 300-450 g (males) or 200-350 g (females) before surgery. We maintained the rats under a reverse 12:12 h light/dark cycle (lights off at 8:00 a.m.) with food and water freely available. We housed two rats/cage prior to surgery and individually after surgery. We excluded 4 rats due to failure to acquire heroin self-administration (n=1, female), faulty catheter on day 8 of training (n=1, male), or data points considered statistical outliers identified using the “Explore” procedure of the “Descriptive Statistics” module of SPSS (n=2, one male and one female). We performed the experiments in accordance with the NIH Guide for the Care and Use of Laboratory Animals (8th edition) under protocols approved by the Animal Care and Use Committee of the Intramural Research Program of the National Institute on Drug Abuse.

### Drugs

We received heroin hydrochloride (HCl) and buprenorphine HCl from the NIDA pharmacy. We dissolved heroin in sterile saline and chose a unit dose of 0.1 and 0.05 mg/kg/infusion for self-administration training based on our previous work (Bossert et al. 2004; Bossert et al. 2020; Bossert et al. 2022). We dissolved buprenorphine in a solution of 20% dimethyl sulfoxide (DMSO), 15% ethanol (EtOH), and sterile water at concentrations that yielded chronic delivery of 6 mg/kg/day dose; the dose of buprenorphine was based on our previous study using chronic buprenorphine delivery (Bossert et al. 2020). The investigators were not blind to the minipump dose conditions.

### Surgery

#### Intravenous surgery

We anesthetized the rats with isoflurane (5% induction; 2-3% maintenance, Covetrus). We attached silastic catheters to a modified 22-gauge cannula cemented to polypropylene mesh (Industrial Netting), inserted the catheter into the jugular vein, and fixed the mesh to the mid-scapular region of the rat (Bossert et al. 2022; Caprioli et al. 2015). We injected the rats with ketoprofen (2.5 mg/kg, s.c., Covetrus) during surgery and on the following day to relieve pain and decrease inflammation. We also injected Enrofloxacin (2.27% diluted 1:9 in sterile saline, s.c., Covetrus) before surgery, 2-3 days post-surgery, and if we observed an infection during the experiment. Rats recovered for 6-8 days before heroin self-administration training. During all experimental phases, we flushed the catheters daily with gentamicin in sterile saline (5 mg/mL, 0.1 mL, Covetrus).

#### Minipump surgery

Chronic delivery of buprenorphine was achieved by implanting osmotic minipumps subcutaneously (Alzet model 2ML2, 5□μl/h for 14-16 days, Durect Corporation) as described in our previous studies (Bossert et al. 2024; Bossert et al. 2020; Bossert et al. 2022). We anesthetized the rats with isoflurane as described above, injected them with ketoprofen, and made a small incision on the right side of the intravenous backmount. We used a hemostat to spread apart the subcutaneous connective tissue to make a small pocket for the pump. We placed the osmotic pumps into the pocket with the flow moderator directed away from the incision. We then closed the incisions with sterile surgical suture and/or wound clips.

### Apparatus

We trained and tested the rats in standard Med Associates self-administration chambers. Each chamber had two levers located 7.5-8 cm above the grid floor on opposing walls. Lever presses on the active, retractable lever activated the infusion pump, whereas lever presses on the inactive, non-retractable lever had no programmed consequences. As in our previous studies (Bossert et al. 2024; Bossert et al. 2019; Bossert et al. 2020; Bossert et al. 2022), the active lever was inserted into the chamber at the start of the self-administration and test sessions and retracted at the end of these sessions, while the inactive lever was constantly present in the chamber. The two contexts differed in their auditory, visual, and tactile cues, as in our previous studies (Bossert et al. 2024; Bossert et al. 2019; Bossert et al. 2020; Bossert et al. 2022). We refer to the contexts as A and B, where A is the context for self-administration training and B is the context for extinction. We counterbalanced the physical environments of contexts A and B.

### Behavioral procedures

The experiment consisted of four phases (Figure 1A): Heroin self-administration training in context A (12 days, 6-h/day), Day 1 extinction test in context B (1 day, 30 min test), minipump implantation surgery (1 day) and Day 21 or 22 extinction test in context B (1 day, 2-h test).

**Figure 1.**
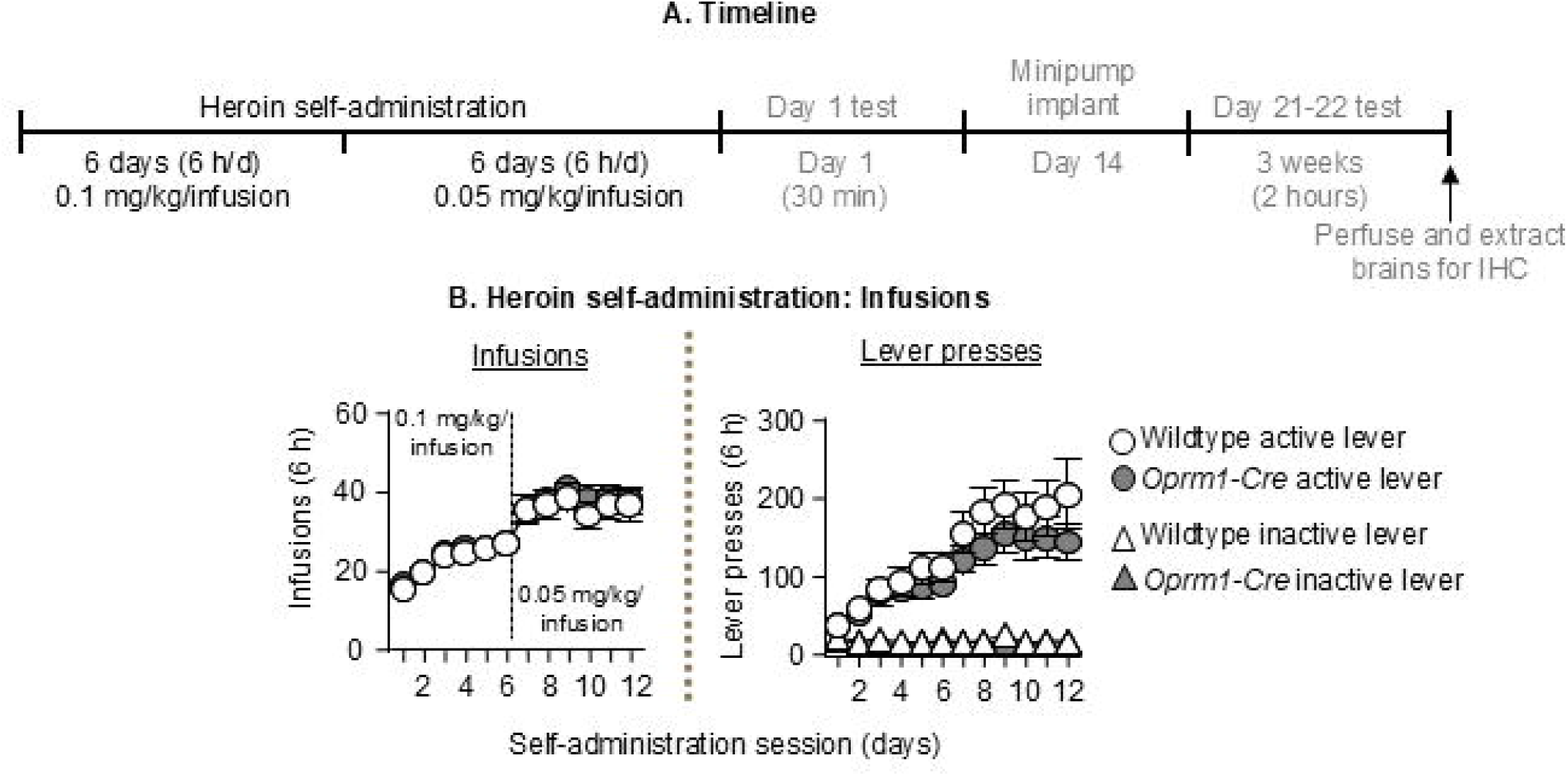
Heroin self-administration training. **(A)** Timeline **(B)** Number of heroin infusions (left panel) and lever presses (right panel) in *Oprm1-Cre* rats and wildtype littermates during heroin self-administration (n=52, 27 *Oprm1-Cre*, 25 wildtype, 20 males, 32 females). All data are mean□ ±□ SEM.

### Heroin self-administration training

We trained the rats to self-administer heroin HCl in context A for 6 h/day (six 1-h sessions separated by 10 min) for 12 days. Each session began with the illumination of a houselight that remained on for the entire session; the active lever was inserted into the chamber 10 s after the houselight was illuminated. During training, the rats earned heroin infusions by pressing on the active lever; infusions were paired with a compound tone–light cue for 3.5 s under a fixed ratio 1 (FR1) reinforcement schedule with a 20-s timeout after each infusion. Heroin was infused at a volume of 100 µl over 3.5 s at a dose of 0.1 mg/kg/infusion (the first 6 sessions) and then 0.05 mg/kg/infusion (the last 6 sessions). Responses on the active lever during the timeout period were recorded but did not result in heroin infusions. Presses on the inactive lever were recorded but had no programmed consequences. At the end of each session, the houselight was turned off and the active lever was retracted. If we suspected catheter failure during training, we tested patency with Diprivan (propofol, NIDA pharmacy, 10 mg/mL, 0.1-0.2 mL injection volume, i.v.). If the catheter was not patent, we catheterized the left jugular vein.

### Tests for heroin seeking on abstinence day 1 and days 21-22

We first tested rats in a brief (30 min) extinction session in context B one day after the last day of heroin self-administration training (abstinence day 1). During the test session, responses on the previously active lever led to presentations of the tone-light cue but not heroin infusions. On abstinence day 14, we implanted the rats with Alzet minipumps containing vehicle or buprenorphine (6 mg/kg/day). We used the first 30 min of the 2-h extinction session on abstinence day 21 or 22 to evaluate whether chronic buprenorphine delivery would decrease incubation of heroin seeking. We tested the rats in context B because our previous study showed reliable incubation of heroin seeking in this context after one week of homecage abstinence (Bossert et al. 2022). We also selected context B for incubation testing because we originally planned to compare, in future studies, the effects of specific causal manipulations in discrete brain regions (e.g., Daun02 inactivation, selective lesions of MOR-expressing cells) on incubation with their effects on context-induced reinstatement of heroin seeking after extinction and on the acquisition of heroin self-administration in context A (Bossert et al. 2020).

There were no genotype differences in heroin training or day 1 extinction responding. We tested wildtype littermates on day 21 and tested *Oprm1-Cre* rats on day 22; we perfused all rats on the same day (day 22). This allowed for us to use the wildtype rats as a baseline (No Test) control for Fos expression against which to evaluate incubation-associated Fos expression in the different brain regions in the *Oprm1-Cre* rats. As mentioned in the introduction, our initial goal was to co-localize Fos+Cre in the *Oprm1-Cre* rats so that we could quantify Fos expression in Cre-labeled (*Oprm1*-positive) cells. However, we were unable to obtain specific Cre labeling despite piloting three different anti-Cre primary antibodies raised in rabbit, mouse, and guinea pig.

### Tissue processing for immunohistochemistry

The Fos and MOR immunohistochemistry procedures is based on our previous work (Bossert et al. 2016; Bossert et al. 2012). Two hours after the relapse (incubation) test on abstinence day 22, we deeply anesthetized the rats with isoflurane saturated in air in an enclosed glass desiccator for approximately 80 s and perfused them transcardially with 100 ml of 0.1M phosphate-buffered saline (PBS) followed by ∼400 ml of 4% paraformaldehyde in 0.1M sodium phosphate (pH 7.4). We removed the brains and post-fixed in 4% paraformaldehyde for 2 h before transferring them to 30% sucrose in 0.1 M sodium phosphate (pH 7.4) for 48 h at 4°C. We subsequently froze the brains in powdered dry ice and stored them at −80°C until sectioning. Using a cryostat (Leica Microsystems), we sectioned the brains (coronal plane) approximately +4.0 mm to +1.0 mm (40 µm) from Bregma in four series, and stored them in PBS containing 0.1% sodium azide at 4°C.

### Double labelling Fos-MOR immunohistochemistry (IHC)

We performed IHC on a subset of rats (n=33 out of 52) from the 4 test conditions (see methods below): No Test Vehicle (n=7), No Test 6 mg/kg/day (n=7), Test Vehicle (n=10), and Test 6 mg/kg/day (n=9). We rinsed free-floating sections three times for 10 min each in PBS, quenched the sections for 15 min with 0.3% hydrogen peroxide (H_2_O_2_) and then rinsed the sections three more times in PBS. We then incubated for 2 h in 4% Bovine Serum Albumin (BSA) in PBS with 0.3% Triton X-100 (PBS-Tx) and incubated overnight at 4°C with rabbit anti-Fos primary antibody (1:8000; Phospho-c-Fos, catalog #5348S, RRID: AB_10557109, Cell Signaling Technology) in 4% BSA in 0.3% PBS-TX. The next day, we rinsed the sections in PBS and incubated for 2 h with biotinylated anti-rabbit IgG secondary antibody (1:600, catalog #BA-1000, RRID: AB_2313606, Vector Labs) in 4% BSA in 0.3% PBS-TX. We rinsed the sections in PBS and incubated in avidin-biotin-peroxidase complex (ABC Elite kit, PK-6100, Vector Labs) in 0.5% PBS-Tx for 1 h. We then rinsed the sections in PBS and performed the reaction for Fos using Vector SG (blue/gray product; Vector SG peroxidase substrate kit, catalog #SK-4700, Vector Labs) for approximately 6 minutes. We then rinsed the sections several times for 5 min each in PBS, incubated in 0.3% H_2_O_2_ for 30 min, and then rinsed three more times in PBS before incubation for 2 h in 0.3% PBS-Tx containing 4% BSA and avidin D (Avidin-Biotin blocking kit, catalog #SP-2001, Vector Labs).

We then incubated the sections overnight at 4°C with rabbit anti-MOR primary antibody (1:1000, catalog # ab134054, RRID:AB_3122135, abcam) (Bailly et al. 2020; Corder et al. 2017; Friedman et al. 2017; Lupp et al. 2011) in 4% BSA, 0.3% PBS-Tx, and biotin (Avidin-Biotin blocking kit). We rinsed the sections with PBS and incubated for 2 h with biotinylated anti-rabbit IgG secondary antibody (1:200) in 4% BSA in 0.3% PBS-TX. We again rinsed the sections in PBS and then incubated in avidin-biotin-peroxidase complex (ABC Elite kit) in 0.5% PBS-Tx for 1 h. We then rinsed the sections in PBS and performed the reaction for MOR in 3,3’-diaminobenzidine (DAB) for approximately 3 minutes. We rinsed the sections in PBS, mounted onto chrom-alum/gelatin-coated slides, and air-dried. We dehydrated the slides through a graded series of alcohol concentrations (30, 60, 90, 95, 100, 100% ethanol), cleared them with Citrasolv (Fisher Scientific), and cover slipped them with Permount (Fisher Scientific).

### Analysis and quantification of Fos and MOR

We captured brightfield images of Fos and MOR immunoreactive (IR) cells using an Olympus VS200 Slide Scanner (10x objective). We quantified cells in mPFC, OFC, and striatum using the following coordinates from Bregma: +3.7 to +3.0 mm for orbitofrontal cortex (OFC), +3.5 to +2.5 mm for anterior cingulate cortex (Cg1), prelimbic cortex (PL), infralimbic cortex (IL), dorsal peduncular cortex (DP), and +2.0 to + 1.2 mm for dorsolateral and dorsomedial striatum (DLS and DMS, respectively), NAc core, and NAc shell. For all brain areas except the striosomes and matrix compartments of the DLS and DMS, we defined regions of interest (ROIs), outlined with pink rectangle boundaries in Figures 3-4, based on the 6^th^ edition of the Paxinos and Watson rat brain atlas (Paxinos and Watson 2007). For DLS and DMS, we quantified Fos in striosomes and matrix separately. We chose 4 striosomes per area and per hemisphere with the densest MOR staining. We defined ROIs of striosomes based on the densest MOR staining, shown by dotted pink boundaries in Figure 4 top DMS and DLS panels. For the matrix, we used similar sized ROIs as that for striosomes to quantify Fos in matrix adjacent to striosomes’ regions without MOR staining. We manually identified cells with high Fos expression (dark brown nuclei) and quantified the number of Fos-positive cells in 2 sections (3-4 hemispheres) and calculated the mean of these counts per area mm^2^ (i.e., each rat provided an individual data point for data analyses with data averaged from 2 images per brain area for each rat). We performed image-based quantification of Fos-positive cells in ImageJ in a blind manner (inter-rater reliability between JMB and HB *Pearson r*=0.87-0.98; RP and HW, *r*=0.99-1.00).

### Statistical analysis

We analyzed the behavioral and immunohistochemistry data with repeated-measures analysis of variance (RM-ANOVA) ANOVAs using SPSS (Version 31.0, GLM procedure). We describe the specific between- and within-subjects factors of each analysis in the Results section. We used both males and females per NIH mandate but combined their data for the statistical analyses because we did not power our study to detect sex differences. We also combined the sexes because previous studies provided little evidence for sex differences in incubation of opioid seeking after forced or voluntary abstinence (Chow et al. 2025; Fredriksson et al. 2021; Nicolas et al. 2022). We used Lever (active, inactive) as a factor in the statistical analyses but do not show the inactive lever data in the figures because during the relapse (incubation) tests, inactive lever presses were very low (2.4±0.4 per 30 min on day 1 and 6.8±1.0 per 2 h on day 21/22) and there were no genotype differences. Finally, because our RM-ANOVAs yielded multiple main and interaction effects, we only report in the Results section significant effects that are critical for data interpretation. For complete statistical results, see Tables S1 and S2.

## Results

### Heroin self-administration training (Fig. 1 and Table S1)

We trained the rats to self-administer heroin at 0.1 mg/kg/infusion for the first 6 days, followed by 0.05 mg/kg/infusion for the next 6 days. Rats of both genotypes showed reliable heroin self-administration as indicated by an increase in the number of heroin infusions and active lever presses over days, and a compensatory increase in the number of infusions earned when we halved the dose (Fig. 1B, see Table S1 for complete statistical). There was no significant genotype difference in heroin self-administration (Table S1). There were also no sex differences across genotypes in heroin self-administration (data not shown).

### Effect of chronic delivery of buprenorphine on incubation of heroin seeking (Fig. 2 and Table S2)

Active lever presses in the vehicle condition were higher on abstinence day 21/22 than on day 1 (incubation of heroin seeking). Chronic delivery of buprenorphine significantly decreased incubation of heroin seeking in both genotypes.

**Figure 2.**
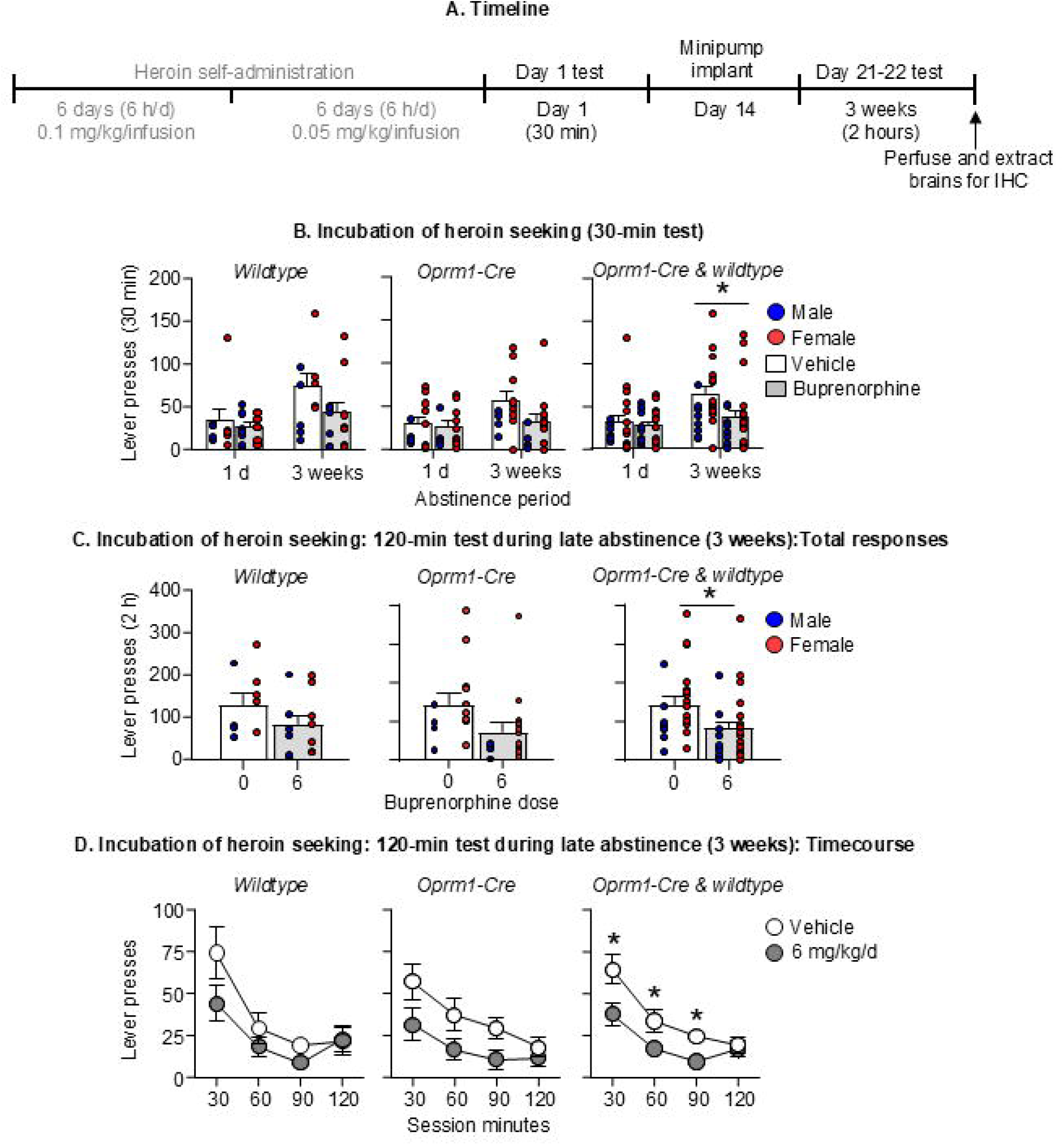
Effect of chronic buprenorphine on incubation of heroin seeking. **(A)** Timeline **(B)** Number of active lever presses on abstinence Day 1 (before minipump implantation) and on abstinence Day 21-22 (first 30 min). During the 30-min extinction sessions, active lever presses led to contingent presentations of the tone-light clue previously paired with heroin, but not heroin. **(C)** Number of active lever presses during the 120-min extinction session and time course during the four 30-min sessions. Vehicle: (n=23; 10 wildtype [5 males, 5 females] 13 *Oprm1-Cre* [4 males, 9 females]). Buprenorphine (6 mg/kg/d): (n=29, 15 wildtype [7 males, 8 females] and 14 *Oprm1-Cre* [4 males, 10 females]). All data are mean □ ± □ SEM. * Different from Vehicle, p<0.05.

The RM-ANOVA of the 30-min lever-pressing data of abstinence day 1 and the first 30-min of the day 21/22 included the between-subjects factors of Genotype (wildtype, *Oprm1-Cre*) and Buprenorphine Dose (0, 6 mg/kg/day), and the within-subjects factors of Day (1, 21-22) and Lever (inactive, active). This analysis showed significant effects of Buprenorphine Dose (F[1,48]=5.3, p=0.025], Day (F[1,48]=14.7, p<0.001], Lever (F[1,48]=95.9, p<0.001], and a significant Day x Buprenorphine dose interaction (F[1,48]=12.6, p=<0.001]. There was no main effect of Genotype or an interaction between the two factors.

The RM-ANOVA of the 2-h lever-pressing data of abstinence day 21/22 included the between-subjects factors of Genotype and Buprenorphine Dose, and the within-subjects factors of Session Time (30, 60, 90, 120 min) and Lever. This analysis showed significant effects of Buprenorphine Dose (F[1,48]=7.0, p=0.011], Session Time (F[3,144]=31.5, p<0.001], Lever (F[1,48]=75.4, p<0.001], and a significant Session Time x Buprenorphine dose interaction (F[3,144]=3.7, p=0.013]. There was no main effect of Genotype or an interaction between the two factors.

Finally, visual inspection of the data (see Fig. 2) suggested that, unlike previous studies (Nicolas et al.2022), incubation of heroin seeking in the vehicle condition was stronger in females than in males. This impression was supported by post-hoc analyses that includes Sex in the RM-ANOVAs, that showed a significant Sex effect for the analyses of both the 30-min data of abstinence day 1 and day 21/22 (F[1,44]=5.5, p=0.024], and the 2-h data of day 21/22 (F[1,44]=5.1, p=0.029]. However, there were no significant interactions of Sex with the other experimental factors.

### Effect of chronic buprenorphine on incubation-associated Fos expression (Fig. 3-4 and Table S3)

**Figure 3.**
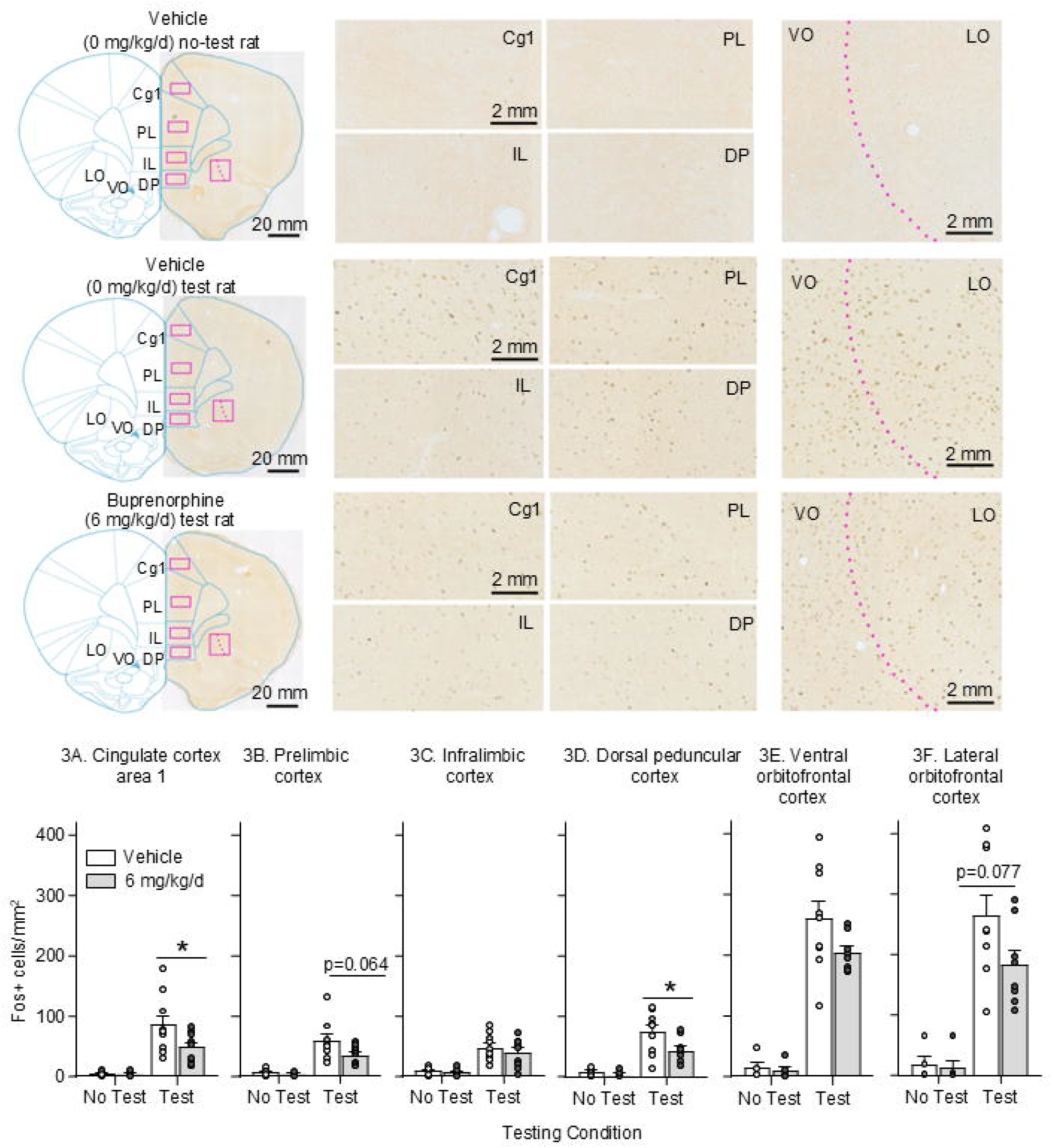
Effect of chronic buprenorphine on Fos expression in PFC. (**A-F**) Representative regions of interest (ROI) locations (pink rectangles) for Fos and images of Fos-IR in mPFC (cingulate cortex 1, prelimbic cortex, infralimbic cortex, and dorsal peduncle cortex) and OFC (ventral and lateral) in rats perfused either immediately after the Day 21-22 test or 24-h later (No test, Test). Atlas templates modified from (Paxinos and Watson 2007). <u>Fos-IR quantification:</u> Number of Fos-positive cells per mm^2^ in the different brain regions. mPFC: No test: n=13 (6 Vehicle, 7 Buprenorphine), Test: n=19 (10 Vehicle, 9 Buprenorphine). OFC: No test: n=12 (6 Vehicle, 6 Buprenorphine), Test n=17 (9 Vehicle, 8 Buprenorphine). Data are mean□±□SEM. * Different from Vehicle, p □ < □ 0.05. Scalebar, 2 mm.

**Figure 4.**
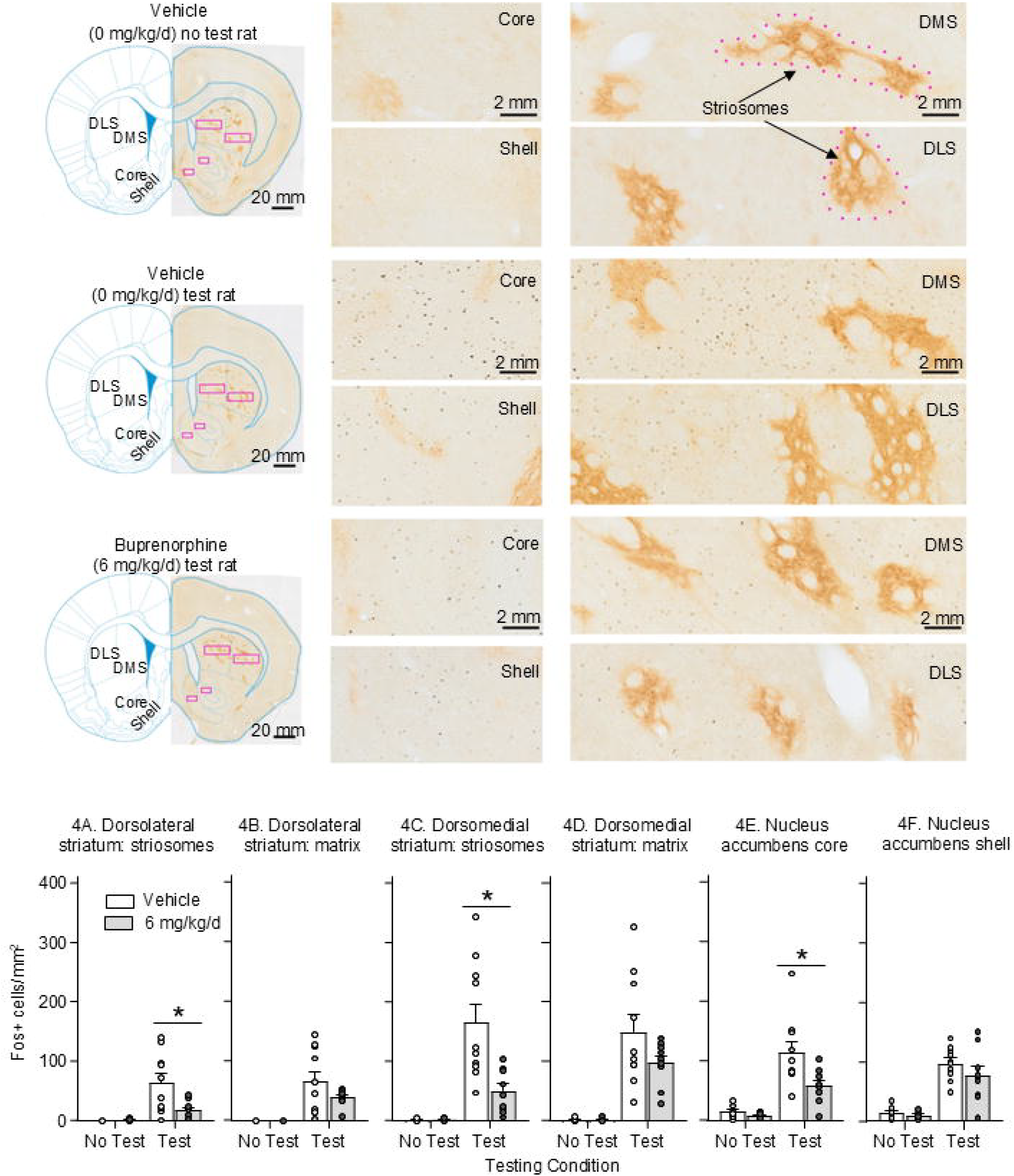
Effect of chronic buprenorphine on Fos expression in striatum. (**A-F**) Representative regions of interest (ROI) locations (pink rectangles) for Fos and images of Fos-IR in dorsal striatum (dorsolateral and dorsomedial striatum, striosomes [examples indicated by dotted pink lines] and matrix of each subregion) and nucleus accumbens (core and shell) in rats perfused either immediately after the Day 21-22 test or 24-h later (No test, Test). Atlas templates modified from (Paxinos and Watson 2007). Fos-IR quantification: Number of Fos-positive cells per mm^2^ in the different brain regions. No test: n=14 (7 Vehicle, 7 Buprenorphine), Test: n=19 (10 Vehicle, 9 Buprenorphine). Data are mean □ ± □ SEM * Different from Vehicle, p □ < □ 0.05. Scalebar, 2 mm.

Incubation of heroin seeking on abstinence day 22 was associated with increased neuronal activation in different mPFC, OFC, dorsal striatum, and NAc subregions, and this effect was decreased by chronic buprenorphine in several subregions.

#### Medial prefrontal cortex (mPFC)

The RM-ANOVA included the between-subjects factors of Test Condition (No Test, Test) and Buprenorphine Dose (0, 6 mg/kg/day) and the within-subjects factor of mPFC subregion (Cg1, PL, IL, and DP). The analysis showed significant effects for Test Condition (F[1,28]=42.6, p<0.001], mPFC subregion (F[3,84]=5.2, p=0.002], mPFC subregion x Test Condition (F[3,84]=7.0, p<0.001], and a marginally significant main effect of Buprenorphine Dose (F[1,28]=3.4, p=0.078]. Post-hoc analyses within each subregion using Test Condition and Buprenorphine dose as the between-subjects factor showed a significant effect of Buprenorphine Dose for DP (F[1,28]=4.7, p=0.039) but not the other subregions (Table S3).

#### Orbitofrontal cortex (OFC)

The RM-ANOVA included the between-subjects factors of Test Condition and Buprenorphine Dose and the within-subjects factor of OFC subregion (ventral, lateral). The analysis showed a significant main effect for Test Condition (F[1, 24]=83.9, p<0.001], but not Buprenorphine Dose or Test Condition x Buprenorphine Dose interaction. (Table S3).

#### Dorsal striatum (DS)

The RM-ANOVA included the between-subjects factors of Test Condition and Buprenorphine Dose and the within-subjects factors of DS subregion (medial, lateral) and DS compartment (striosomes, matrix). The analysis showed significant effects for Test Condition (F[1,29]=30.5, p<0.001], DS subregion (F[1,29]=32.1, p<0.001], DS compartment (F[1,29]=4.4, p=0.046], DS subregion x Test Condition (F[1,29]=29.2, p<0.001), Compartment x Test Condition (F[1,29]=5.0, p=0.033), Compartment x Buprenorphine Dose (F[1,29=10.5, p=0.003], Compartment x Test Condition x Buprenorphine Dose (F[1,29]=11.3, p=0.002), DS subregion x Compartment x Buprenorphine Dose (F[1,29]=14.1, p<0.001), and DS subregion x Compartment x Test Condition x Buprenorphine Dose (F[1,29]=12.1, p=0.002); the effect of Buprenorphine Dose was marginally significant (F[1,29]=4.2, p=0.051). Post-hoc analyses within each subregion and compartment using Test Condition and Buprenorphine dose as the between-subjects factor showed a significant effect of Buprenorphine Dose for DLS and DMS striosomes (F[1,29]=4.6 and 7.6, p=0.041 and 0.010, respectively) and for DLS and DMS striosomes for Test Condition x Buprenorphine Dose (F[1,29]=5.3 and 7.8, p=0.029 and 0.009, respectively) (Table S4).

#### Nucleus accumbens (NAc)

The RM-ANOVA included the between-subjects factors of Test Condition and Buprenorphine Dose and the within-subjects factor of NAc subregion (Core, Shell). The analysis showed significant main effects for Test Condition (F[1,29]=55.1, p<0.001] and Buprenorphine Dose (F[1,29]=4.6, p=0.04]. Post-hoc analyses within each subregion using Test Condition and Buprenorphine Dose as the between-subjects factor showed a significant effect of Buprenorphine Dose (F[1,29=6.0, p=0.021) for NAc Core but not Shell (Table S4).

## Discussion

Buprenorphine is an FDA-approved medication for opioid addiction (Jasinski et al. 1978; Kakko et al. 2003), yet the brain regions mediating its therapeutic effects are unknown. As an initial step, we examined whether buprenorphine-induced inhibition of incubation of heroin seeking is associated with decreased neuronal activation in cortical and striatal regions expressing MOR.

Incubation of heroin seeking on abstinence day 22 was associated with increased neuronal activation in 5/12 tested cortico-striatal regions. In mPFC, chronic buprenorphine decreased incubation-related neuronal activation in anterior cingulate and dorsal peduncular, but not prelimbic or infralimbic subregions. In OFC, buprenorphine did not decrease activation in either ventral or lateral subregion. In striatum, buprenorphine decreased activation in dorsolateral, dorsomedial, and NAc core, but not NAc shell. Within dorsal striatum, the effect of buprenorphine was stronger in MOR-rich striosomes than in the surrounding matrix, where MOR expression is low (Graybiel 1990).

At the behavioral level, the inhibitory effect of buprenorphine on incubation of heroin seeking extends our prior findings on its effects on other relapse-related behaviors (extinction responding, context-induced reinstatement, and reacquisition) in rats trained to self-administer oxycodone (Bossert et al. 2020).

An unexpected result was stronger incubation of heroin seeking in females compared to males under vehicle conditions (see Results). This contrasts with prior studies of incubation of heroin and oxycodone seeking after forced or voluntary abstinence, where no sex differences were observed (Fredriksson et al. 2020; Fredriksson et al. 2021; Venniro et al. 2019; Venniro et al. 2017). Possible explanations include the use of *Oprm1-Cre* knock-in rats and their wild-type littermates (bred in our facility), as opposed to commercially available Sprague-Dawley rats used in most prior studies (Chow et al. 2025). Another possibility is that unlike most previous studies where tests were performed in the self-administration context, we performed the incubation tests in the present study in a novel context (context B). However, this seems less likely, as a prior study in Sprague-Dawley rats did not show sex differences in incubation of heroin seeking when rats were tested in context B after 1 and 8 abstinence days (Bossert et al. 2022).

Several limitations should be considered. First, our study is correlational. Without causal manipulations to inhibit Fos-expressing neurons, it remains unclear whether the observed activation patterns during late abstinence reflects a cause or a consequence of heroin seeking. Additionally, it is unknown whether buprenorphine’s inhibition of neuronal activity reflects a causal mechanism for its behavioral effects on incubation of heroin seeking or simply a pharmacological suppression of neuronal activity. Similarly, the strong inhibition of activation in MOR-rich striosomes may not reflect a causal role for these cells in incubation. Indeed, we recently found that lesions of MOR-expressing dorsal striatum neurons using a Cre-dependent taCasp3 vector in Oprm1-Cre rats (Bossert et al. 2023) did not alter incubation of heroin seeking in the absence of buprenorphine (Bossert JM, unpublished observations).

The contributions of other MOR-expressing cortical and striatal subregions where buprenorphine reduced Fos remain to be determined.

### Concluding remarks

In our initial investigation of brain regions involved in buprenorphine’s inhibitory effects on incubation of heroin seeking, we found that chronic buprenorphine decreased incubation-related brain activation induction in several cortical and striatal subregions. These findings highlight potential brain targets for follow-up mechanistic studies using approaches such as the Daun02 inactivation method (Bossert et al. 2011; Koya et al. 2009) or *Oprm1-Cre* rats (Bossert et al. 2023) to selectively manipulate Fos-positive or MOR-expressing neurons.

Finally form a clinical perspective, our observation that buprenorphine reduces the incubation of heroin seeking provides support for the predictive (or more accurately, postdictive validity (Epstein et al. 2006)) of the incubation of heroin craving model. This point is timely, given recent challenges to the face validity of the model, because clinical studies have not reported time-dependent increases in self-reported heroin craving during abstinence (Bergeria et al. 2024). For a discussion of possible reasons for this discrepancy between animal and human findings, see (Chow et al. 2025).

## Supporting information

SOM

## Ethics declaration

We performed the experiments in accordance with the NIH Guide for the Care and Use of Laboratory Animals (8th edition) under protocols approved by the Animal Care and Use Committee of the Intramural Research Program of the National Institute on Drug Abuse.

## Author’s contributions

JMB, KEC, and MSP performed the surgeries, JMB and KEC ran the experiments and performed the histology. JMB and RM captured and preprocessed the images for analysis. JMB, HB, RP, and HW quantified Fos. JMB and YS designed the study and performed the data analyses. JMB, RM, HW, and YS wrote the manuscript. All authors critically reviewed the content and approved the final version before submission.

## Supplemental Online Material

Table S1 Statistical reporting of heroin self-administration training

Table S2 Statistical reporting of incubation of heroin seeking

Table S3 Statistical reporting of mPFC and OFC Fos quantification

Table S4 Statistical reporting of dorsal and ventral striatum Fos quantification

