## Supplementary material for "Chronic delivery of buprenorphine during abstinence decreases incubation of heroin seeking and neuronal activation in medial prefrontal cortex and striatum in male and female rats": SOM

9-20-2025

**Table of content**

Table S1. Statistical reporting of heroin self-administration training

Table S2. Statistical reporting of incubation of heroin seeking

Table S3. Statistical reporting of mPFC and OFC Fos quantification

Table S4. Statistical reporting of dorsal and ventral striatum Fos quantification

**Table S1.** *Heroin self-administration training.* Statistical analysis (SPSS GLM repeated-measures module). Partial Eta^2^ = proportion of explained variance. RM, repeated measures.

| **Figure number** | **Factor name** | **F-value** | ***p*-value** | **Partial Eta^2^** |
| --- | --- | --- | --- | --- |
| Figure 1B. Heroin self-administration training  Infusions  RM-ANOVA | Genotype (WT, *Oprm1-Cre*) between-subjects  Heroin Dose (0.1, 0.05 mg/kg/infusion) within-subjects  Heroin Dose x Genotype  Session (1-6, 7-12) within-subjects  Session x Genotype  Heroin Dose x Session  Heroin Dose x Session x Genotype | F(1,50)=0.1  F(1,50)=102.3  F(1,50)=0.2  F(5,240)=13.0  F(5,240)=0.4  F(5,240)=7.0  F(5,240)=0.2 | 0.720  <0.001*  0.629  <0.001*  0.856  <0.001*  0.974 | 0.003  0.672  0.005  0.206  0.008  0.122  0.003 |
| Figure 1B. Heroin self-administration training  Lever presses  RM-ANOVA | Genotype (WT, *Oprm1-Cre*) between-subjects  Lever (inactive, active) within-subjects  Lever x Genotype  Heroin Dose (0.1, 0.05 mg/kg/infusion) within-subjects  Heroin x Genotype  Session (1-6, 7-12) within-subjects  Session x Genotype  Lever x Heroin Dose  Lever x Heroin Dose x Genotype  Lever x Session  Lever x Session x Genotype  Heroin Dose x Session  Heroin Dose x Session x Genotype  Lever x Heroin Dose x Session  Lever x Heroin Dose x Session x Genotype | F(1,50)=1.0  F(1,50)=64.2  F(1,50)=0.9  F(1,50)=47.1  F(1,50)=1.5  F(5,240)=9.0  F(5,240)=0.5  F(1,50)=49.7  F(1,50)=1.5  F(5,250)=10.2  F(5,250)=0.7  F(5,250)=2.3  F(5,250)=0.3  F(5,250)=2.6  F(5,250)=0.3 | 0.313  <0.001*  0.345  <0.001*  0.225  <0.001*  0.771  <0.001*  0.231  <0.001*  0.655  0.044*  0.923  0.026*  0.905 | 0.020  0.562  0.018  0.485  0.029  0.153  0.010  0.499  0.029  0.170  0.013  0.044  0.006  0.049  0.006 |

**Table S2.** *Effect of chronic buprenorphine on incubation of heroin seeking.* Statistical analysis (SPSS GLM repeated-measures module). Partial Eta^2^ = proportion of explained variance. RM, repeated measures.

| **Figure number** | **Factor name** | **F-value** | ***p*-value** | **Partial Eta^2^** |
| --- | --- | --- | --- | --- |
| Figure 2B. Day 1 and Day 21/22: 30-min data  Lever presses  RM-ANOVA | Genotype (WT, *Oprm1-Cre*) between-subjects  Buprenorphine Dose (0, 6 mg/kg/d) between-subjects  Genotype x Buprenorphine Dose  Day (1, 21/22) within-subjects  Day x Genotype  Day x Buprenorphine Dose  Day x Genotype x Buprenorphine Dose  Lever (inactive, active) within-subjects  Lever x Genotype  Lever x Buprenorphine Dose  Lever x Genotype x Buprenorphine Dose  Day x Lever  Day x Lever x Genotype  Day x Lever x Buprenorphine Dose  Day x Lever x Genotype x Buprenorphine Dose | F(1,48)=1.6  F(1,48)=5.3  F(1,48)=0.1  F(1,48)=14.7  F(1,48)=0.9  F(1,48)=4.8  F(1,48)=0.0  F(1,48)=95.9  F(1,48)=0.9  F(1,48)=3.6  F(1,48)=0.0  F(1,48)=13.0  F(1,48)=1.0  F(1,48)=2.9  F(1,48)=0.0 | 0.219  0.025*  0.714  <0.001*  0.347  0.033*  0.992  <0.001*  0.345  0.065  0.830  <0.001*  0.319  0.097  0.997 | 0.031  0.100  0.003  0.234  0.018  0.091  0.000  0.666  0.019  0.069  0.001  0.214  0.021  0.056  0.000 |
| Figure 2C. Day 21-22: Total responses (120 min)  Lever presses  RM-ANOVA | Genotype (WT, *Oprm1-Cre*) between-subjects  Buprenorphine Dose (0, 6 mg/kg/d) between-subjects  Genotype x Buprenorphine Dose  Lever (inactive, active) within-subjects  Lever x Genotype  Lever x Buprenorphine Dose  Lever x Genotype x Buprenorphine Dose | F(1,48)=0.3  F(1,48)=7.0  F(1,48)=0.2  F(1,48)=75.4  F(1,48)=0.1  F(1,48)=5.1  F(1,48)=0.2 | 0.567  0.011*  0.668  <0.001*  0.723  0.028*  0.678 | 0.007  0.127  0.004  0.611  0.003  0.096  0.004 |
| Figure 2C. Day 21-22: 30-min time course  Lever presses  RM-ANOVA | Genotype (WT, *Oprm1-Cre*) between-subjects  Buprenorphine Dose (0, 6 mg/kg/d) between-subjects  Genotype x Buprenorphine Dose  Lever (inactive, active)  Lever x Genotype  Lever x Buprenorphine Dose  Lever x Genotype x Buprenorphine Dose  Session Time (30, 60, 120, 180) within-subjects  Session Time x Genotype  Session Time x Buprenorphine Dose  Session Time x Genotype x Buprenorphine Dose  Lever x Session Time  Lever x Session Time x Genotype  Lever x Session Time x Buprenorphine Dose  Lever x Session Time x Genotype x Buprenorphine Dose | F(1,48)=0.3  F(1,48)=7.0  F(1,48)=0.2  F(1,48)=75.4  F(1,48)=0.1  F(1,48)=5.1  F(1,48)=0.2  F(3,144)=31.5  F(3,144)=2.6  F(3,144)=3.7  F(3,144)=0.4  F(3,144)=24.6  F(3,144)=2.3  F(3,144)=2.2  F(3,144)=0.3 | 0.567  0.011*  0.668  <0.001*  0.723  0.028*  0.678  <0.001*  0.054  0.013*  0.788  <0.001*  0.076  0.087  0.856 | 0.007  0.127  0.004  0.611  0.003  0.096  0.004  0.397  0.052  0.072  0.007  0.339  0.047  0.044  0.005 |

**Table S3.** *Effect of chronic buprenorphine on Day 21-22 Fos expression: mPFC and OFC.* Anterior cingulate cortex, Cg1; dorsal peduncle, DP; infralimbic cortex, IL; medial prefrontal cortex, mPFC; prelimbic cortex, PL; OFC, orbitofrontal cortex.

| **Figure number** | **Factor name** | **F-value** | ***p*-value** | **Partial Eta^2^** |
| --- | --- | --- | --- | --- |
| Figure 3. Fos/mm2  **mPFC**  RM-ANOVA and ANOVA subregions | Test Condition (No test, Test) between-subjects  Buprenorphine Dose (0, 6 mg/kg/d) between-subjects  Test Condition x Buprenorphine Dose  mPFC subregion (Cg1, PL, IL, DP) within-subjects  mPFC subregion x Test Condition  mPFC subregion x Buprenorphine Dose  mPFC subregion x Test Condition x Buprenorphine Dose  Cg1: Test Condition  Cg1: Buprenorphine Dose  Cg1: Test Condition x Buprenorphine Dose  PL: Test Condition  PL: Buprenorphine Dose  PL: Test Condition x Buprenorphine Dose  IL: Test Condition  IL: Buprenorphine Dose  IL: Test Condition x Buprenorphine Dose  DP: Test Condition  DP: Buprenorphine Dose  IL: Test Condition x Buprenorphine Dose | F(1,28)=42.6  F(1,28)=3.4  F(1,28)=2.4  F(3,84)=5.2  F(3,84)=7.0  F(3,84)=1.9  F(3,84)=2.0  F(1,28)=34.1  F(1,28)=3.0  F(1,28)=3.0  F(1,28)=32.0  F(1,28)=3.0  F(1,28)=2.2  F(1,28)=29.5  F(1,28)=0.6  F(1,28)=0.3  F(1,28)=42.6  F(1,28)=4.7  F(1,28)=2.8 | <0.001*  0.078  0.129  0.002*  <0.001*  0.131  0.119  <0.001*  0.094  0.095  <0.001*  0.093  0.151  <0.001*  0.455  0.618  <0.001*  0.039*  0.108 | 0.603  0.106  0.080  0.158  0.200  0.065  0.067  0.549  0.097  0.097  0.533  0.097  0.072  0.513  0.020  0.009  0.603  0.144  0.090 |
| Figure 3. Fos/mm2  **OFC**  RM-ANOVA and ANOVA subregions | Test Condition (No test, Test) between-subjects  Buprenorphine Dose (0, 6 mg/kg/d) between-subjects  Test Condition x Buprenorphine Dose  OFC subregion (Ventral, Lateral) within-subjects  OFC subregion x Test Condition  OFC subregion x Buprenorphine Dose  OFC subregion x Test Condition x Buprenorphine Dose  Ventral: Test Condition  Ventral: Buprenorphine Dose  Ventral: Test Condition x Buprenorphine Dose  Lateral: Test Condition  Lateral: Buprenorphine Dose  Lateral: Test Condition x Buprenorphine Dose | F(1,24)=83.9  F(1,24)=2.5  F(1,24)=1.8  F(1,24)=0.1  F(1,24)=0.7  F(1,24)=0.7  F(1,24)=0.6  F(1,24)=111.3  F(1,24)=2.1  F(1,24)=1.5  F(1,24)=55.9  F(1,24)=2.4  F(1,24)=1.9 | <0.001*  0.127  0.187  0.715  0.404  0.402  0.431  <0.001*  0.158  0.231  <0.001*  0.131  0.186 | 0.778  0.094  0.071  0.006  0.029  0.029  0.026  0.823  0.081  0.059  0.700  0.093  0.072 |

**Table S4.** *Effect of chronic buprenorphine on Day 21/22 Fos expression: dorsal striatum and nucleus accumbens***.** Dorsal striatum, DS; dorsolateral striatum matrix, DLS matrix; dorsolateral striatum striosomes, DLS striosomes; dorsomedial striatum matrix, DMS matrix; dorsomedial striatum striosomes, DMS striosomes

nucleus accumbens core, NAc core; nucleus accumbens shell, NAc shell

| Figure 4. Fos/mm2  **DS subregion & Compartment**  RM-ANOVA and ANOVA subregions | Test Condition (No test, Test) between-subjects  Buprenorphine Dose (0, 6 mg/kg/d) between-subjects  Test Condition x Buprenorphine Dose  DS subregion (DLS, DMS) within-subjects  DS subregion x Test Condition  DS subregion x Buprenorphine Dose  DS subregion x Test Condition x Buprenorphine Dose  Compartment (Striosomes, Matrix) within-subjects  Compartment x Test Condition  Compartment x Buprenorphine Dose  Compartment x Test Condition x Buprenorphine Dose  DS subregion x Compartment  DS subregion x Compartment x Test Condition  DS subregion x Compartment x Buprenorphine Dose  DS subregion x Compartment x Test Condition x Buprenorphine Dose  DLS matrix: Test Condition  DLS matrix: Buprenorphine Dose  DLS matrix: Test Condition x Buprenorphine Dose  DMS matrix: Test Condition  DMS matrix: Buprenorphine Dose  DMS matrix: Test Condition x Buprenorphine Dose  DLS striosomes: Test Condition  DLS striosomes: Buprenorphine Dose  DLS striosomes: Test Condition x Buprenorphine Dose  DMS striosomes: Test Condition  DMS striosomes: Buprenorphine Dose  DMS striosomes Test Condition x Buprenorphine Dose | F(1,29)=30.5  F(1,29)=4.2  F(1,29)=4.3  F(1,29)=32.1  F(1,29)=29.2  F(1,29)=3.7  F(1,29)=3.6  F(1,29)=4.4  F(1,29)=5.0  F(1,29)=10.5  F(1,29)=11.3  F(1,29)=0.6  F(1,29)=0.3  F(1,29)=14.1  F(1,29)=12.1  F(1,29)=22.9  F(1,29)=1.4  F(1,29)=1.4  F(1,29)=38.4  F(1,29)=1.7  F(1,29)=1.8  F(1,29)=15.4  F(1,29)=4.6  F(1,29)=5.3  F(1,29)=25.8  F(1,29)=7.6  F(1,29)=7.8 | <0.001*  0.051  0.046*  <0.001*  <0.001*  0.063  0.067  0.046*  0.033*  0.003*  0.002*  0.456  0.593  <0.001*  0.002*  <0.001*  0.240  0.240  <0.001*  0.202  0.194  <0.001*  0.041*  0.029*  <0.001*  0.010*  0.009* | 0.512  0.125  0.130  0.525  0.502  0.114  0.111  0.130  0.147  0.265  0.281  0.019  0.010  0.326  0.294  0.442  0.047  0.047  0.570  0.055  0.057  0.347  0.136  0.154  0.471  0.209  0.211 |
| --- | --- | --- | --- | --- |
| Figure 4. Fos/mm2  **NAc subregion**  RM-ANOVA and ANOVA subregions | Test Condition (No test, Test) between-subjects  Buprenorphine Dose (0, 6 mg/kg/d) between-subjects  Test Condition x Buprenorphine Dose  NAc subregion (Core, Shell) within-subjects  NAc subregion x Test Condition  NAc subregion x Buprenorphine Dose  NAc subregion x Test Condition x Buprenorphine Dose  Core: Test Condition  Core: Buprenorphine Dose  Core: Test Condition x Buprenorphine Dose  Shell: Test Condition  Shell: Buprenorphine Dose  Shell: Test Condition x Buprenorphine | F(1,29)=55.1  F(1,29)=4.6  F(1,29)=2.6  F(1,29)=0.0  F(1,29)=0.0  F(1,29)=2.8  F(1,29)=2.3  F(1,29)=35.1  F(1,29)=6.0  F(1,29)=3.8  F(1,29)=53.9  F(1,29)=1.5  F(1,29)=0.6 | <0.001*  0.040*  0.115  0.853  0.985  0.106  0.140  <0.001*  0.021*  0.060  <0.001*  0.231  0.435 | 0.655  0.137  0.084  0.001  0.000  0.088  0.074  0.548  0.170  0.117  0.650  0.049  0.021 |
